# Ambient temperature effects on stress-induced hyperthermia in Svalbard ptarmigan

**DOI:** 10.1101/594614

**Authors:** Andreas Nord, Lars P. Folkow

## Abstract

Stress-induced hyperthermia (SIH) is commonly observed during handling in homeotherms. However, in birds, handling in cold environments typically elicits hypothermia. It is unclear whether this indicates that SIH is differently regulated in this taxon or if it is due to size, because body temperatures changes during handling in low temperature have only been measured in small birds ≤ 0.03 kg (that are more likely to suffer high heat loss when handled). We have, therefore, studied thermal responses to handling stress in the intermediate-sized (0.5-1.0 kg) Svalbard ptarmigan (*Lagopus muta hyperborea*) in 0°C and −20°C, in winter and spring. Handling caused elevated core body temperature, and peripheral vasoconstriction that reduced back skin temperature. Core temperature increased less and back skin temperature decreased more in −20°C than in 0°C, probably because of higher heat loss rate at the lower temperature. Responses were qualitatively consistent between seasons, despite higher body condition/insulation in winter and dramatic seasonal changes in photoperiod, possibly affecting stress responsiveness. Our study supports the notion that SIH is a general thermoregulatory reaction to acute stressors in endotherms, but also suggests that body size and thermal environment should be taken into account when evaluating this response in birds.

## INTRODUCTION

Stress-induced hyperthermia (SIH), is a ubiquitous feature of the body’s response to acute stressors, such as restraint or an altered social context, in mammals and birds (e.g. Briese and Cabanac, 1991; Cabanac and Briese, 1992; Carere and Van Oers, 2004; Gray et al., 2008; Korhonen et al., 2000; Meyer et al., 2008). It is believed that the increase in core body temperature (*T*_c_) during SIH represents a sympathetically mediated elevation of the hypothalamic set point, i.e. an ‘active hyperthermia’ resembling pathogen-induced fevers (Briese and Cabanac, 1991; Kluger et al., 1987; Oka et al., 2001; Vinkers et al., 2009). Hence, SIH is sometimes referred to as ‘stress fever’ or ‘psychogenic fever’ (e.g. IUPS Thermal Commission, 2003), although several studies suggest it employs different neural pathways (Gray et al., 2008; Vinkers et al., 2009). Thermal responses leading to elevated *T_c_* during SIH include cutaneous vasoconstriction and shivering thermogenesis (e.g. Briese and Cabanac, 1991; Herborn et al. 2015; Jerem et al., 2015; Kluger et al., 1987; Oka et al., 2001). The diversion of peripheral blood flow to the core, together with stress-induced tachycardia and increased ventilation rate (Cabanac and Aizawa, 2000; Cabanac and Guillemette, 2001; Greenacre and Lusby, 2004; Mans et al., 2012), probably prepares the animal for escape or interaction (i.e., a “fight or flight” response), at the same time as centralization of the blood pool and increased blood clotting function could minimize blood loss in the event of injury (Cannon 1915).

Because SIH probably reflects set point change it is predicted that, for a given stressor, *T*_c_ changes should be independent of the ambient temperature (*T*_a_) under which stress is perceived. This prediction is supported by laboratory studies of rodents (e.g. Briese and Cabanac, 1991; Kluger et al., 1987; Long et al., 1990). However, studies of birds have revealed remarkable variation in the thermal responses to restraint or handling at different *T*_a_’s. Specifically, handling in thermoneutral conditions sometimes invokes SIH (Cabanac and Aizawa, 2000; Cabanac and Guillemette, 2001; Carere and Van Oers, 2004; Herborn et al., 2015), sometimes hypothermia (Maggini et al., 2018; Møller, 2010), and sometimes seem to leave *T*_c_ unaltered (Lewden et al., 2017), whereas handling below thermoneutrality consistently seems to elicit hypothermia without any co-occurring signs of shock such as lack of muscle tonus (Andreasson et al., 2019; Lewden et al., 2017; Udvardy, 1955;).

Cooling rather than warming of the core need not mean that birds have differentially regulated stress responses to handling. Rather, because the cooling phenomenon seems confined to small (< 0.03 kg) species with inherently higher thermal conductance, it is likely that stress-associated *hypothermia* results from changes to the rate and avenues of heat transfer caused by the handling *per se*. For example, a small bird enclosed by a hand is subject to a substantial increase in the proportion of surface area amenable for conductive heat transfer, suffers reduced insulation when the plumage is compressed, and probably conducts heat over a considerably steeper thermal gradient (as a consequence of the handling-induced reduction in plumage depth). This explanation remains speculative, though, because body temperature responses to handling in larger, better insulated, birds have never been recorded in low *T*_a_ (e.g. Cabanac and Aizawa, 2000; Cabanac and Guillemette, 2001; Herborn et al., 2015). In addition, there are no continuous data from birds on the relationship between temperature changes in the periphery and the core during stress, which further complicates our understanding of the regulatory processes involved in situations where ‘atypical’ thermal responses are observed. Thus, additional data from large and well-insulated birds measured in low *T*_a_ are required to better understand why this taxon displays such variable stress-induced body temperature responses.

We measured the thermal responses to handling in the Svalbard ptarmigan (*Lagopus muta hyperborea* Sundevall) (Fig. 1). This 0.5 to 1.0 kg bird is endemic to the high-arctic Svalbard archipelago (76-81°N), where it is exposed to *T*_a_ ranging ca. −40 to +20°C and a photoperiod varying 24 h over the course of the year. Its thermal conductance varies accordingly, being the lowest in winter when it is dark and cold and birds are in prime body condition, and the highest in summer when birds are at their leanest and have moulted into a less insulating plumage (Mortensen and Blix 1986; Nord & Folkow 2018). These properties make the Svalbard ptarmigan a good model for studying how body temperatures of a relatively large bird respond to handling, and if these responses differ in different thermal environments and in line with variation in insulation. Accordingly, we measured temperature changes both peripherally (back and head skin; assumed to reflect changes in cutaneous circulation) and in the body core (to assess whether birds responded with hypo- or hyperthermia) in Svalbard ptarmigan subjected to handling stress at thermoneutrality (0°C) and far below thermoneutrality (−20°C) (Mortensen and Blix, 1986), both in winter when insulation peaked, and in spring when insulation was declining. Measurement of both deep and peripheral temperatures, and at times of year when thermal conductance is notably different (Mortensen and Blix, 1986; Nord and Folkow, 2018), was expected to provide information on the thermoregulatory processes involved in acute stress in birds. If the normal thermal response to stress in birds is an increase in *T*_c_ (i.e., SIH), and the reduced *T*_c_ during handling in smaller species of birds is, in fact, a consequence of increased heat loss rate (e.g. from plumage disturbances or increased conductive cooling; cf. Andreasson et al., 2019; Lewden et al., 2017) rather than representing a differentially regulated response, we predicted that the larger and well-insulated Svalbard ptarmigan would show an elevation of *T*_c_ that was qualitatively similar at thermoneutral (0°C) vs. very low (−20°C) *T*_a_’s, and that this would be preceded by reduced peripheral temperature (reflecting cutaneous vasoconstriction). Because the Svalbard ptarmigan has low thermal conductance compared to other species, even at its leanest in summer (Mortensen & Blix 1986), we also predict that any seasonal effect on the thermal responses to handling should be minor.

**Fig. 1.**
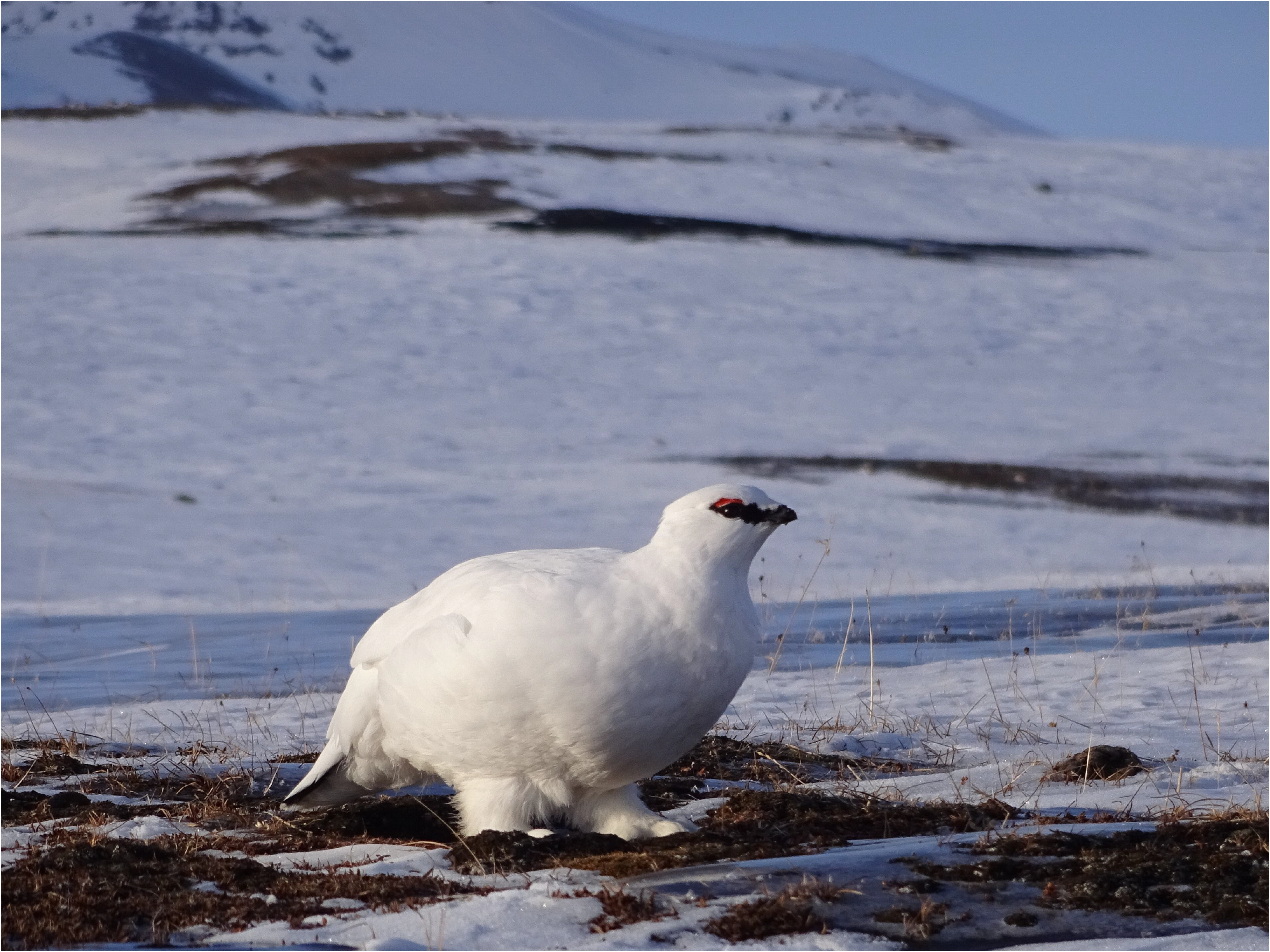
Male Svalbard ptarmigan in winter plumage overlooking the slopes of the Arctowski mountain, Spitsbergen, Svalbard (78°12’ N, 16°17’ E) in early spring. Photo © Andreas Nord

## MATERIALS AND METHODS

### Birds and housing

Twelve male Svalbard ptarmigan were used. Six were captured as chicks near Longyearbyen, Svalbard (78°13’ N, 15°38’ E) 16 to 17 months before the start of the experiment, and the other six were captive bred at the University of Tromsø – the Arctic University of Norway. At the time of the study, all females in the facility were part of the breeding population that is exempt from experimental work year round. However, in Svalbard ptarmigan, both sexes show the same seasonal responses and are of similar size (Mortensen et al., 1983), suggesting that stress responsiveness should be qualitatively similar in males and females. All birds were maintained indoors in thermoneutrality under simulated Longyearbyen photoperiod. The morphological and physiological changes associated with winter acclimation do not differ between captive and wild-caught Svalbard ptarmigan as long as the birds are maintained in natural photoperiod (Lindgård and Stokkan, 1989; Stokkan et al., 1986). Housing conditions followed Nord and Folkow (2018).

### Measurement of body temperature and handling protocol

All birds were physically mature (i.e., older than 1 year). Studies were undertaken under polar night conditions in early winter (DD) (07 December 2015 to 17 January 2016), when body condition peaked (body mass: 1064.2 ± 33.0 g (s.d.); dissectible fat: 261.5 ± 16.7 g) and in spring (03 May to 03 June 2016) in continuous light (LL) when body condition was declining (body mass: 828.1 ± 18.1 g; dissectible fat: 142.0 ± 9.2 g). Data were collected at ambient temperatures (*T*_a_) 0°C (*T*_a_ = −0.1 ± 0.6°C (s.d.), which is within the thermoneutral zone, and at −20°C (−19.9 ± 0.5°C), which is far below thermoneutrality in Svalbard ptarmigan independent of season (Mortensen and Blix, 1986; Nord and Folkow, 2018). We used 11 birds in winter, of which 10 were measured in 0°C and five also in −20°C. One additional bird was measured only in −20°C. Seven of the 11 winter birds, and one additional bird, were measured in spring (i.e., 0°C: *n* = 8 birds; −20°C: *n* = 5 birds, of which 4 had been measured in −20°C also in winter). In winter, 7 birds were first measured in 0°C, and 4 birds were first measured in −20°C. In spring, all birds were first measured in 0°C and then in −20°C. Repeated measurement within seasons were spaced 27 ± 3 (s.d.) days (range 24 to 31 days). Birds were monitored as part of a study that is not reported here for 21 days after each measurement session. For this reason, sampling in both *T*_a_’s in winter could not be achieved within the period of constant darkness. In spring, measurement in both *T*_a_’s was not always possible, since some individuals were allocated to the on-site breeding program by the end of May.

Starting at 10:06 am ± 30 min (s.d.) (GMT +1), birds were weighed and fitted with 36-gauge type T (copper-constantan) thermocouples (Omega Engineering, Stamford, CT, USA) for measurement of *T*_c_ (70 mm into the colon), back skin temperature (*T*_back_) between the wings, and head skin temperature (*T*_head_) on the scalp following Nord & Folkow (2018), within 10-15 min of collection from the cage. Skin thermocouples were attached using a small amount of cyanoacrylate glue (Loctite^®^ Power Easy Gel, Henkel, Düsseldorf, Germany) and taking care not to cover the thermocouple junction. This glue does not cause any skin damage or irritation in Svalbard ptarmigan (AN, LPF, pers. obs.). Thermocouples were calibrated at 0°C (Ice point drywell model 5115) and 40°C (High precision bath model 6025, both Fluke Calibration, American Fork, UT, USA) prior to use.

Once thermocouples were attached and before the handling experiment, we placed the bird in a 33.6 l transparent acrylic glass metabolic chamber (that was ventilated with ambient air at 5.1 ± 0.3 l min^−1^) inside a climatic chamber (model 24/50 DU, Weiss Technik, Giessen, Germany) for measurement of resting metabolic rate and body temperatures during 1 h 44 min ± 13 min (s.d.), as part of a different study. Both metabolic rate and body temperatures stabilized at lower levels within 30 min after instrumentation. After completion of metabolic measurements, we opened the metabolic chamber, side-pinned (terminology *sensu* Herborn et al., 2015) the bird, and administered an immune challenge (100 μl 1 mg kg^−1^ intramuscular LPS), also as part of the other study that is not reported here. Instead, we here report the changes in body temperatures that were recorded during and after the handling stress that was induced thereby. This stressor lasted, on average, 3.2 ± 0.6 min (s.d.) (range: 2.5 to 5.3 min) and did not differ in length between seasons or *T*_a_’s (season × *T*_a_, season, *T*_a_: all *P* ≥ 0.3). Thus, the order of events during measurements were: capture and instrumentation (10-15 min); equilibration and baseline data collection at relevant *T*_a_ (104 min); handling stress (3 min) and post-stressor data collection (20 min). Because the metabolic chamber was fully open when birds were handled, metabolic rate could not be measured during, and in the 20 min period following, stress exposure.

During winter measurements, the climatic chamber was always dark (except for illumination with dim red light, ≪ 1 lx, to allow video inspection) to simulate polar night. In spring, the chamber was always fully illuminated by full spectrum white light bulbs. All but three birds had been subjected to a similar handling- and measurement protocol on 2 to 8 instances in a previous study (mean ± s.d.: 6 ± 2) (for details on this protocol, see Nord and Folkow, 2018). Data were recorded at 10 samples/s, and were digitized from raw signals using a ML796 PowerLab/16SP A-D converter (ADInstruments, Sydney, Australia).

Ethical approval for experimental procedures and reuse of birds was issued by the Norwegian Food Safety Authority (permit no. 6640). Live capture and import of Svalbard ptarmigan chicks was under permissions issued by the Governor of Svalbard (permit no. 2014/00290-2 a.522-01), the Norwegian Environment Agency (permit 2018/7288) and the Norwegian Food Safety Authority (permit no. 2014/150134).

### Data analyses

We only used data from periods where the birds were at full rest (standing or walking but not moving vigorously, or perched with ptiloerection) immediately before handling. This criterion was met in 24 of 31 instances (winter, 0°C: 8 of 10; winter, −20°C: 5 of 6; spring, 0°C: 7 of 8; spring, −20°C: 3 of 5). We also dismissed data from *T*_head_ thermocouples that fell off or broke (winter, −20°C: 1; spring, 0°C: 2; spring, −20°C: 1). *T*_c_ and *T*_back_ data were complete.

Pre-handling data were collected during 2 min immediately before handling (i.e., ≥ 1.5 h after the start of body temperature measurement). Handling and post-handling data were then collected during 20 min after the end of this stressor, i.e., a period sufficient to encompass the body temperature response to acute stress in other gallinaceous birds (Herborn et al., 2015), but not long enough to include any thermal or metabolic effects of LPS (Marais et al., 2011). We binned data in 5 s (i.e., 50 samples) averages, partitioned in pre-handling, handling, and post-handling, periods. We then calculated: ‘response amplitude’ as the maximum or minimum body temperature attained during handling; ‘duration of response’ as the first time point where a 30 s running mean for each body temperature had returned to, or intersected, pre-handling values once the stressor had been removed; and ‘response magnitude’ as mean body temperature change (relative to pre-handling values) over the duration of the response. If a given body temperature had not returned to pre-handling levels by the end of the focal period, we set ‘duration of response’ to 20 min (i.e. the length of the post-handling observation period), and calculated ‘response magnitude’ based on this period.

Statistics were done using R 3.4.3 for Windows (R Core Team, 2018). We analysed all response variables in tissue-specific mixed effect models fitted with maximum likelihood (lmer function in the lme4 package) (Bates et al., 2015), with season and *T*_a_ and their interaction as factors, and ‘bird id’ as a random intercept to account for repeated measurements. We did not include body mass or body condition in any models, because both parameters varied strongly between, but only very little within, seasons (above). These metrics, therefore, conveyed largely the same statistical information as ‘season’, so testing for any of the former in presence of the latter was not warranted. Final models were derived by sequentially excluding terms with the highest *P*-values based on likelihood ratio tests for the original and alternative models, starting with the interactions, until only significant (*P* ≤ 0.05) variables remained. We then refitted the final model using restricted maximum likelihood (Zuur et al., 2009) and calculated predicted means ± s.e.m. using the emmeans package (Lenth, 2019).

## RESULTS

Parameter estimates and test statistics are reported in Table S1.

All *T*_c_ responses to handling were positive and attenuated in −20°C compared to in 0°C. Accordingly, birds reached 0.19 ± 0.06°C greater maximum *T*_c_ in 0°C (Δ *T*_c_ = 0.52 ± 0.04°C) than in −20°C (Δ *T*_c_ = 0.32 ± 0.05°C) (*P* = 0.009, Fig. 2). Response duration, i.e. the time taken for *T*_c_ to return to pre-handling levels for at least 30 s once the stressor had been removed, was longer in 0°C (13.4 ± 1.1 min) than in −20°C (5.6 ± 1.4 min) by 7.8 ± 1.8 min (+138%) (*P* < 0.001, Fig. 2). The response magnitude, i.e. the mean deviation from pre-handling *T*_c_ during the response, was larger when birds were measured in thermoneutrality (0.29 ± 0.03°C) than when they were measured in −20°C (0.17 ± 0.04°C), by 0.12 ± 0.04°C (*P* = 0.016). The modifying effect of *T*_a_ was always uniform across seasons (season × *T*_a_: all *P*≥ 0.22), and there was no difference in mean responses between winter and spring (*P* ≥ 0.43 in all cases; Table S1).

**Fig. 2.**
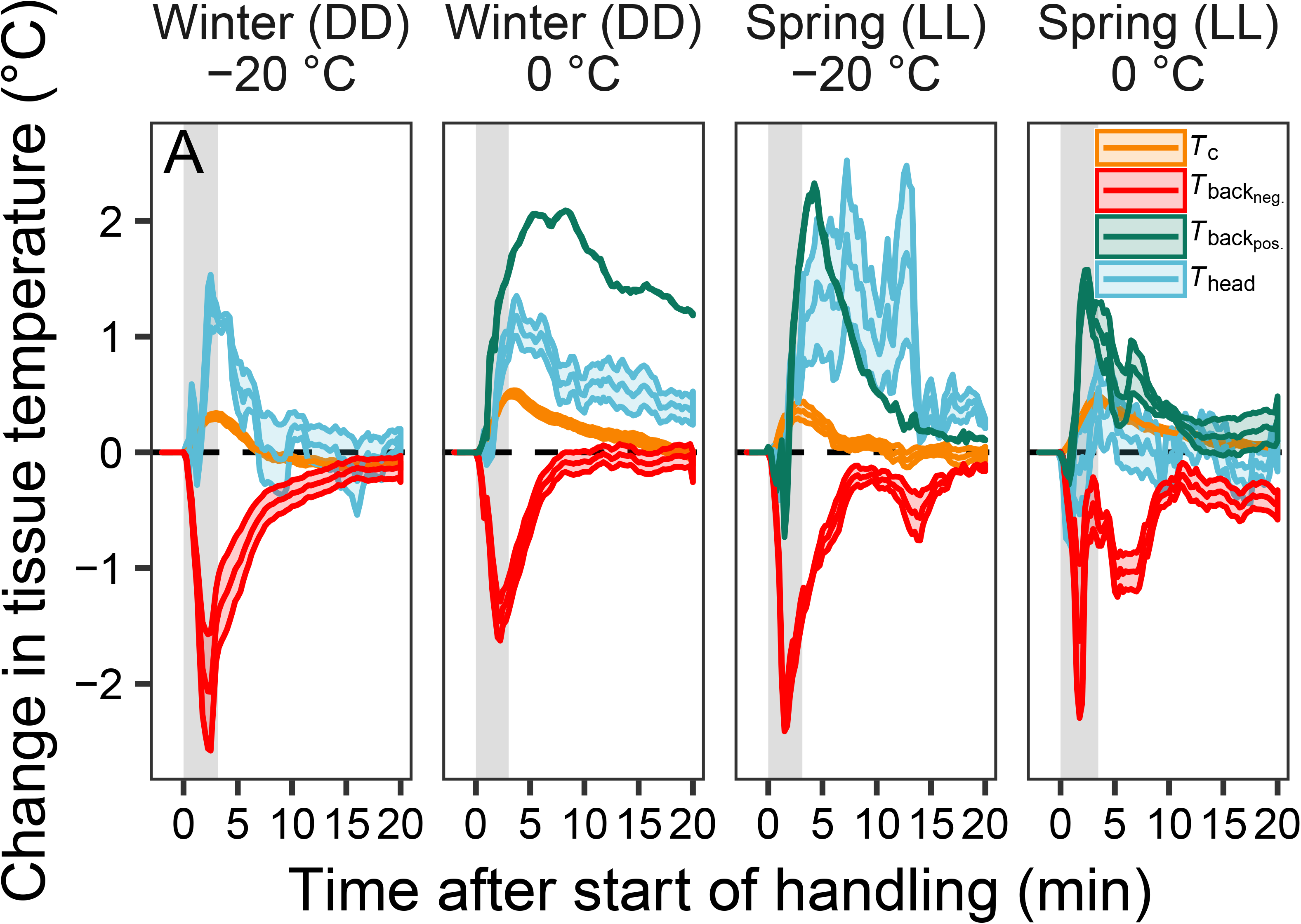
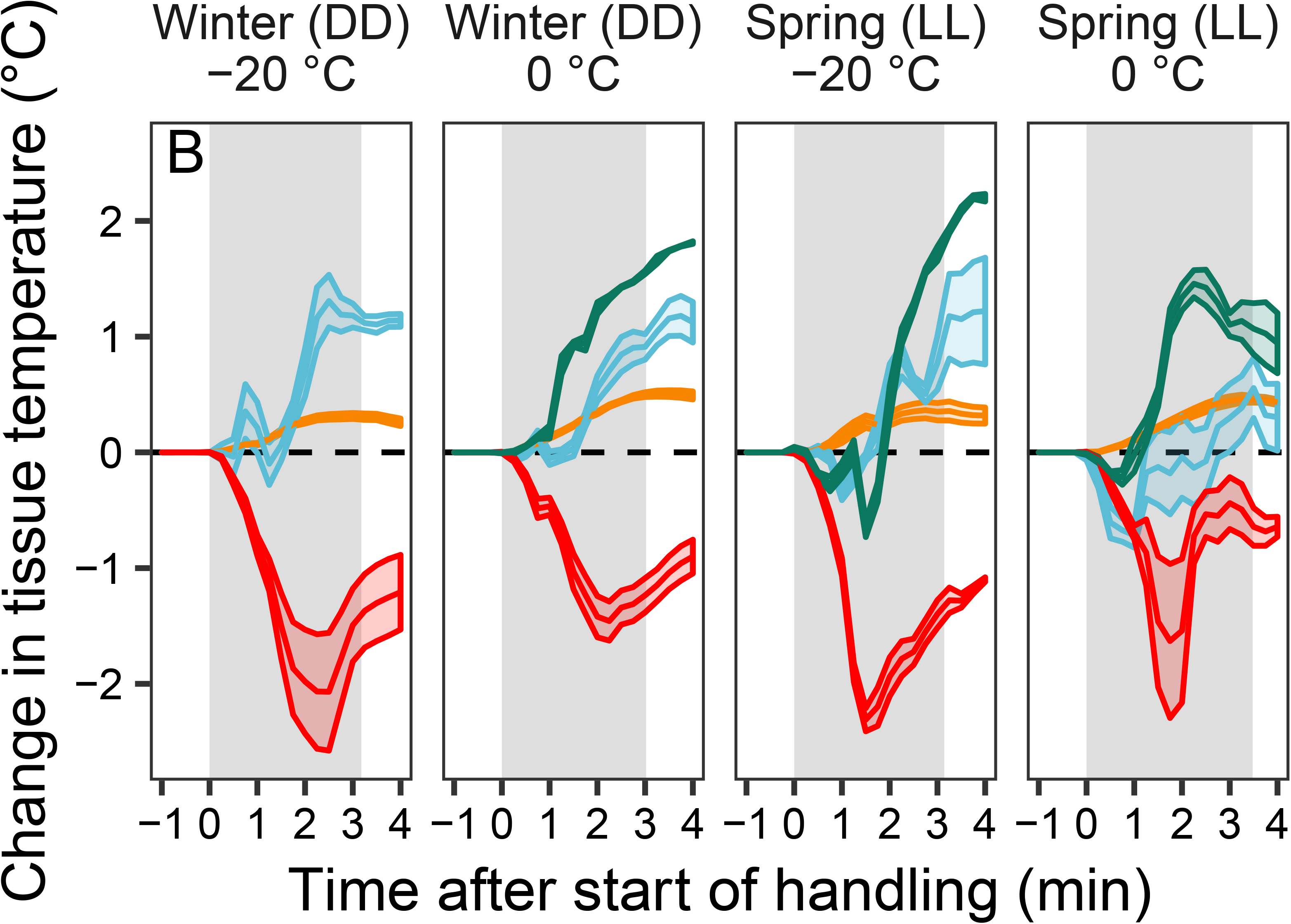
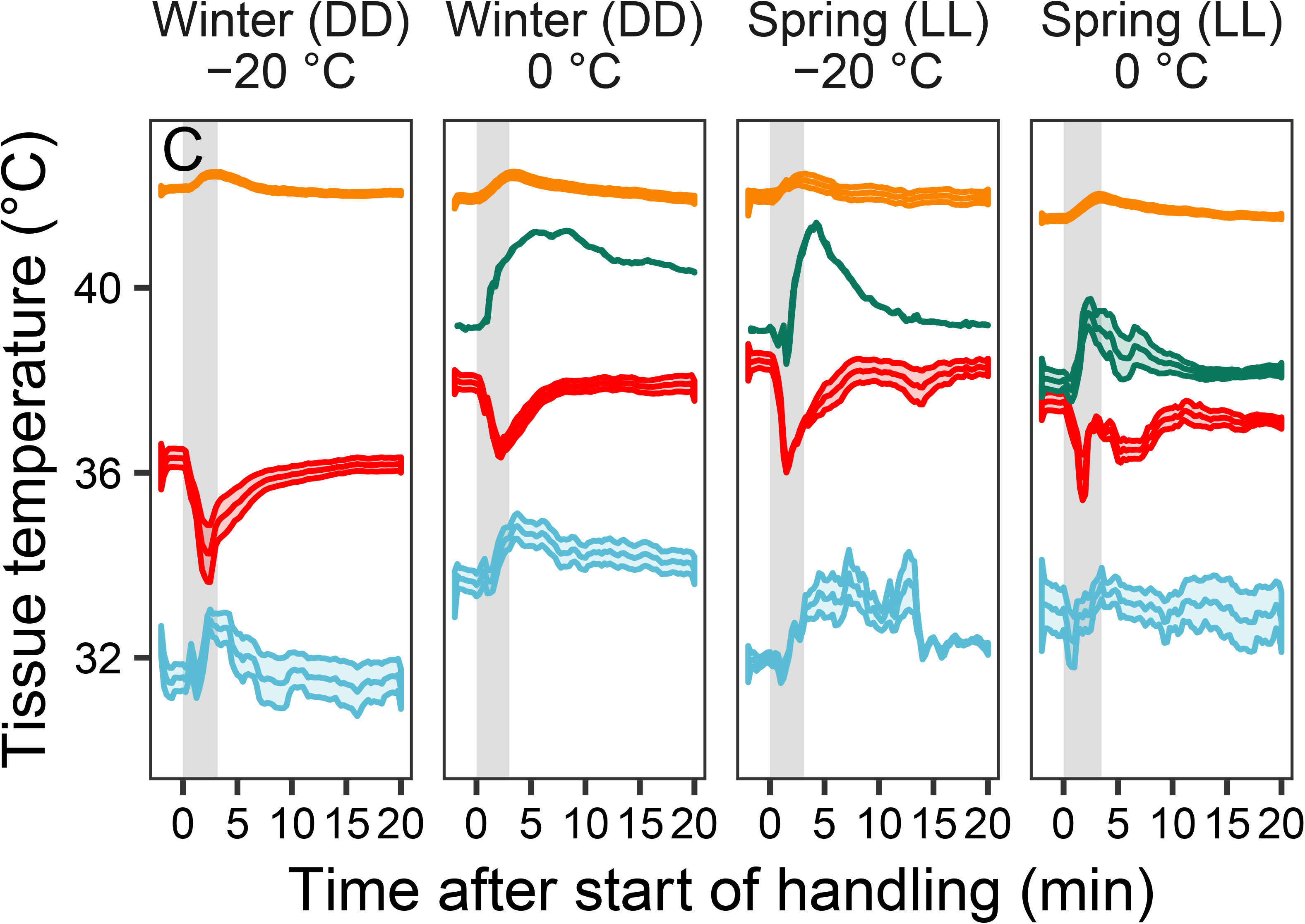
Mean ± s.e.m. changes in cloacal temperature (*T*_c_ – orange), back skin temperature (*T*_back_ – red or green), and head skin temperature (*T*_head_–blue) during handling in Svalbard ptarmigan. Panels show the response over: (A) the 22 min observation period relative to pre-handling temperature; (B) the response during the handling event ± 1 min relative to pre-handling temperature; and (C) mean ± s.e.m. absolute body temperatures during the observation period. Data were collected in thermoneutrality (0°C) and below thermoneutrality (−20°C) in continuous darkness (DD) in winter (−20°C: *n* = 6; 0°C: *n* = 10) and continuous light (LL) in spring (−20° C: *n* = 5; 0°C: *n* = 8). The vertical grey bars show periods of handling. *T*_back_ changes are presented separately for birds showing negative (red) and positive (green) averages responses, of which the latter occurred in three birds in three different measurement conditions. Data were binned in 30 s (A, C) or 15 s (B) intervals before plotting. Metrics extracted from the depicted thermal responses, and ditto analyses, are described in Materials & Methods.

*T*_back_ decreased on average during handling. However, one bird showed increased *T*_back_ in three out of four measurement sessions, and two additional birds showed positive response when measured in thermoneutrality in spring (Fig 1). In all but one of these cases, positive responses were preceded by a slight *T*_back_ decrease (Fig. 2B). The strongest Δ*T*_back_ response was 1.38 ± 0.44°C below pre-handling levels. There was a tendency for a stronger maximum response at −20°C (−2.03 ± 0.77°C) than at 0°C (−1.01 ± 0.53°C) (*P* = 0.092; Fig. 2). Birds had recovered this decrease within 10.5 ± 1.3 min, but tended to take longer to do so at −20°C (12.0 ± 2.2 min) than at 0°C (7.2 ± 1.8 min) (*P* = 0.098; Fig. 2). Average response magnitude was uniform, at 0.45 ± 0.16°C below pre-handling *T*_back_, across *T*_a_’s. Neither season nor the season-by-*T*_a_ interaction affected the *T*_back_ response to handling (all *P* ≥ 0.12; Table S1).

The maximum Δ*T*_head_ response was positive on average (+0.67 ± 0.35°C) (Fig. 2). This increase subsided within 7.5 ± 0.4 min after removal of the stressor (Fig. 2). The response magnitude index showed that Δ*T*_head_ was elevated, on average by 0.45 ± 0.16°C, during this time. Season, *T*_a_, and their interaction, did not affect any *T*_head_ metric (all *P* ≥ 0.1; Table S1).

## DISCUSSION

Handling of Svalbard ptarmigan was associated with increased *T*_c_ and *T*_head_ and decreased *T*_back_ that lasted, on average, 6.6 to 9.7 min after the stressor had been removed (Fig. 2). The maximum *T*_c_ response was small but distinct and within the range of those recorded over similar time periods in other bird species of comparable body sizes but at higher *T*_a_ (Bittencourt et al., 2015; Cabanac and Aizawa, 2000; Cabanac and Guillemette, 2001; Gray et al., 2008). Both the maximum increase in *T*_c_ and the *T*_c_ response magnitude were more pronounced in 0°C than in −20°C (Fig. 2; Table S1). This probably reflects that low *T*_a_ blunted the rate of increase in *T*_c_. Thus, since the stressor was of fixed duration and probably too short to allow the birds to reach a new set-point, *T*_c_ did not have time to increase as much in −20°C as it did in 0°C. *T*_back_, moreover, tended to decrease about twice as much in −20°C than in 0°C (Fig. 2). This could suggest that a stronger peripheral vasoconstrictor response was employed to elevate *T*_c_ when *T*_a_ was lower, but more likely reflects that the drop in skin temperature when cutaneous blood flow was diminished in the cold, was larger in the lower *T*_a_ due to a more rapid heat loss rate. Thus, the difference in response strength in the two *T*_a_’s most likely reflected that higher heat loss rate in the cold slowed and blunted the *T*_c_ response, and increased *T*_back_ change, relative to changes in the milder *T*_a_.

In some cases (5 of 24), birds responded to handling by vasodilation of back skin (reflected by increased *T*_back_; Fig. 2), but this was preceded by an initial *T*_back_ decrease in all but one of these instances. This suggests that also those few birds that displayed positive *T*_back_ change at least initially met the stressful stimulus with the expected cutaneous vasoconstrictor response, but then, for some reason, rapidly switched to vasodilation. Our data do not allow us to determine the proximate explanation for this result, which could be related to skin thermocouple placement in relation to vascular structures, or that the thermal state of these particular birds caused them to revert to heat dissipation because a new set-point temperature was reached more rapidly. In this latter context, we noted that pre-handling *T*_back_ was higher (by 1.05°C) in these birds.

*T*_head_ on the scalp increased in response to handling, but only after a short-lasting drop that implied rapid vasoconstriction (Fig. 2). Thus, cutaneous *T*_head_ initially followed predictions for peripheral temperature change during acute stress, but rapidly reverted to the opposite response. The temperature increase was probably a combined effect of increased delivery of internal heat to the head in conjunction with the rise in *T*_c_, and increased blood flow to the brain in response to the stressor (Hasler et al., 2007; Wang et al., 2005) that together outweighed any reduced supply of heat due to constriction of head skin vasculature. Increased blood delivery to the brain likely prompted higher attentiveness and increased cognitive ability that would aid the animal in decision-making in a threatening situation. Physiologically, increased temperature in the poorly insulated head (cf. Nord and Folkow, 2018), lasting several minutes after removal of the stressor, probably contributed to reversal of the *T*_c_ response due to its effects on central thermoreceptors, even if the ultimate explanation for *T*_head_ change in our study was not related to heat dissipation as such. It, therefore, appears that this response, together with birds’ general heat loss rate, was sufficient to regain thermal balance in the relatively low *T*_a_’s applied in this study. This could also explain why we mostly did not record any corresponding compensatory peripheral temperature increase in back skin as *T*_c_ returned to normothermia (Fig. 2B). This is in contrast to previous studies of the body temperature responses to handling conducted in considerably milder *T*_a_, e.g. in the cold-tolerant common eider (*Somateria mollissima*) (Cabanac and Guillemette, 2001).

## CONCLUSIONS

The Svalbard ptarmigan, like other medium-sized to large birds, displayed a rapid cutaneous vasoconstrictor response and an increase in *T*_c_ during handling, indicative of SIH. These responses did not vary qualitatively between seasons despite higher overall heat loss rate in spring (Nord and Folkow, 2018) that should allow more rapid return to pre-handling *T*_c_. Moreover, even when insulation was close to its annual minimum in late spring, handling in Ta far below thermoneutrality did not cause hypothermic *T*_c_. This is in stark contrast to data from the considerably smaller cold-tolerant black-capped chickadee (*Poecile atricapillus*) (Lewden et al., 2017) and great tit (*Parus major*) (Andreasson et al., 2019), in which handling in the cold results in *T*_c_ dropping several degrees C below normothermic values. Thus, our data from cold exposure in the substantially larger Svalbard ptarmigan support the argument that stress-related *hypothermia* in small birds handled below thermoneutrality probably does not reflect a regulated process but more likely is due to increased heat loss rate following mechanical distortion of insulation during handling, or reduced heat production if handling elicits tonic immobility (cf. Hohtola, 1981). Yet, thermal responses to handling in our birds were still modified by *T*_a_, particularly in the core but to some extent also peripherally on the back. Hence, our study suggests that when interpreting the body temperature responses to handling, one must take into account both body size of the model and the thermal environment in which measurements are undertaken.

## ACKNOWLEDGEMENTS

Hans Lian, Hans Arne Solvang and Renate Thorvaldsen excellently cared for birds outside experimental periods. Comments from Katherine Herborn and Paul Jerem improved a previous version of the manuscript.

## AUTHOR CONTRIBUTION

AN conceived the idea, AN and LPF developed methodology and AN collected the data. AN analysed the data and wrote the first draft, which was critically edited by LPF. Both authors approved the final version of the manuscript and agreed to be accountable for all contents. AN administered the project and acquired the funding.

## DATA AVAILABILITY

Data are deposited in figshare (doi: 10.6084/m9.figshare.8080934).

