## Supplementary material for "Ambient temperature effects on stress-induced hyperthermia in Svalbard ptarmigan": Table S1

**Table S1.** Parameter estimates, likelihood ratios (LRT), and *P*-values (single term deletions), for final models, and *P*-values and test statistics for excluded terms, for models describing body temperature responses to handling in Svalbard ptarmigan in thermoneutrality (0°C) and below thermoneutrality (-20°C) in constant darkness in winter and constant light in spring.

| **Parameter** | **Estimate (SE)** | **LRT** | ***P*** |
|---|---|---|---|
| **Core body temperature ($T_c$)** | | | |
| *Response amplitude(°C)* | | | |
| Final model: | | | |
| $T_a$ (-20°C or 0°C) | | 6.82 | **0.009** |
| $T_a$ = -20°C | 0.32 (0.05) | | |
| $T_a$ = 0°C | 0.52 (0.04) | | |
| Dropped terms: | | | |
| Season (Winter or Spring) | | 0.00 | 0.976 |
| Season × $T_a$ | | 0.28 | 0.596 |
| *Response duration (min)* | | | |
| Final model: | | | |
| $T_a$ (-20°C or 0°C) | | 14.62 | **< 0.001** |
| $T_a$ = -20°C | 5.65 (1.43) | | |
| $T_a$ = 0°C | 13.43 (1.11) | | |
| Dropped terms: | | | |
| Season (Winter or Spring) | | 0.62 | 0.431 |
| Season × $T_a$ | | 0.90 | 0.343 |
| *Response magnitude (°C)* | | | |
| Final model: | | | |
| $T_a$ (-20°C or 0°C) | | 5.81 | **0.016** |
| $T_a$ = -20°C | 0.17 (0.04) | | |
| $T_a$ = 0°C | 0.29 (0.03) | | |
| Dropped terms: | | | |
| Season (Winter or Spring) | | 0.03 | 0.868 |
| Season × $T_a$ | | 0.18 | 0.672 |
| **Back skin temperature ($T_{back}$)** | | | |
| *Response amplitude(°C)* | | | |
| Final model: | | | |
| - | | | |
| Dropped terms: | | | |
| $T_a$ (-20°C or 0°C) | | 2.85 | 0.092 |
| Season (Winter or Spring) | | 1.77 | 0.183 |
| Season × $T_a$ | | 1.06 | 0.303 |
| *Response duration (min)* | | | |
| Final model: | | | |
| - | | | |
| Dropped terms: | | | |
| $T_a$ (-20°C or 0°C) | | 2.74 | 0.098 |
| Season (Winter or Spring) | | 0.02 | 0.881 |
| Season × $T_a$ | | 1.37 | 0.242 |
| *Response magnitude (°C)* | | | |
| Final model: | | | |
| - | | | |
| Dropped terms: | | | |
| Season (Winter or Spring) | | 2.39 | 0.122 |
| $T_a$ (-20°C or 0°C) | | 1.60 | 0.205 |
| Season × $T_a$ | | 0.58 | 0.447 |
| **Head skin temperature ($T_{head}$)** | | | |
| *Response amplitude(°C)* | | | |
| Final model: | | | |
| - | | | |
| Dropped terms: | | | |
| Season (Winter or Spring) | | 0.52 | 0.472 |
| $T_a$ (-20°C or 0°C) | | 0.58 | 0.448 |
| Season × $T_a$ | | 1.40 | 0.236 |
| *Response duration (min)* | | | |
| Final model: | | | |
| - | | | |

| | | |
|---|---|---|
| Dropped terms: | | |
| Season (Winter or Spring) | 0.02 | 0.891 |
| $T_a$ (-20°C or 0°C) | 0.00 | 0.946 |
| Season × $T_a$ | 0.37 | 0.543 |
| *Response magnitude (*°C*)* | | |
| Final model: | | |
| - | | |
| Dropped terms: | | |
| Season (Winter or Spring) | 0.34 | 0.560 |
| $T_a$ (-20°C or 0°C) | 0.07 | 0.785 |
| Season × $T_a$ | 2.21 | 0.137 |
